# Addressing the Southeast Asian snaring crisis: impact of 11 years of snare removal in a biodiversity hotspot

**DOI:** 10.1101/2023.01.26.525728

**Authors:** Andrew Tilker, Jürgen Niedballa, Hung Luong Viet, Jesse F. Abrams, Lucile Marescot, Nicholas Wilkinson, Benjamin M. Rawson, Rahel Sollmann, Andreas Wilting

## Abstract

Unsustainable snaring is causing biodiversity declines across tropical protected areas, resulting in species extinctions and jeopardizing the health of forest ecosystems. Here, we used 11 years of ranger collected data to assess the impact of intensive snare removal on snaring levels in two protected areas in Viet Nam. Snare removal resulted in significant declines in snare occupancy (36.9, 95% BCI [4.6, 59.0] reduction in percent area occupied), but snaring levels nonetheless remained high (31.4, [23.6, 40.8] percent area occupied), and came with a substantial financial cost. Our results indicate that snare removal remains an important component of efforts to protect tropical protected areas, but by itself, is likely insufficient to address this threat. To stop snaring in protected areas, a multifaceted approach will be necessary that combines short-term reactive snare removal with long-term proactive programs that address the underlying drivers behind snaring.

## Introduction

Unsustainable hunting has defaunated tropical forests around the world (Harrison et al, 2016; Benítez-López et al, 2017, 2019), and preventing further defaunation is a priority for addressing global biodiversity loss (Laurance et al 2012; Barlow et al 2018). Furthermore, because large vertebrates perform ecological functions that are integral to maintaining healthy tropical ecosystems (Lacher et al, 2019), the loss of forest fauna jeopardizes a number of ecological services linked to human well-being (Krause & Tilker et al, 2022). For example, defaunation has been linked to reduced carbon storage capacities in tropical forests (Bello et al, 2015; Osuri et al, 2016), increase in prevalence of zoonotic diseases (Young et al, 2014), and loss of ethnocultural identity for indigenous peoples (Bogoni et al, 2020).

Despite an increase in recent years in protected area coverage and implementation of site-based conservation strategies (Watson et al, 2014), mammal and bird populations have undergone severe declines in some regions (Benítez-López et al, 2017, 2019). One of the main causes of wildlife declines in tropical forests is through the use of nonselective snares. Snares are cheap, easy to set in large numbers, and highly effective at capturing terrestrial vertebrate species (Gray et al, 2021). Recent studies have documented snare-driven defaunation as a significant threat to ground-dwelling mammals and birds across Africa (Fa & Brown, 2009; Bi et al, 2017) and Asia (Campbell et al, 2019; Belecky & Gray, 2020; Groenenberg et al, 2023). Snaring has been particularly severe in Southeast Asia, where it has depressed wildlife populations in many protected areas, and remains a significant and ongoing threat (Belecky & Gray, 2020; Gray et al, 2021). A recent study, for example, found that snaring is a more immediate and severe threat to Southeast Asian faunal communities than forest degradation in some areas (Tilker & Abrams et al, 2019). To prevent further defaunation in Southeast Asia, and thus maintain the ecosystem services and functions of its tropical forests, it is important to assess the effectiveness of conservation actions designed to counteract the ongoing ‘snaring crisis’.

Although several conservation actions have been proposed to counter snaring – including general demand reduction and shifting of consumer preferences (Wilkie & Carpenter, 1999), legislative reform combined with adequate prosecution and conviction of offenders (Gray et al, 2021), and supporting informal community guardianship mechanisms to maximize deterrence (Viollaz et al, 2022) – snare removal remains the primary strategy employed in Southeast Asia (Belecky & Gray, 2020). Snare removal is popular because it is straightforward and non-controversial compared to other responses such as arrest and prosecution (Belecky & Gray, 2020), and it will likely continue to prevail as the main approach to address snaring across most protected areas. Some studies have assessed the impact of snare removal on snaring levels, with both positive (Linkie et al, 2015) and equivocal (Becker et al, 2013) findings. Most of these studies, however, have been over relatively short time periods or in areas where snaring levels do not reach the high levels reported for many parts of Southeast Asia (Jachmann, 2008; Becker et al, 2013; Watson et al, 2013). The extent to which snare removal reduces snaring pressure over longer time horizons remains unexplored, hindering an objective evaluation of its effectiveness. For protected areas operating under limited resources, this lack of knowledge is worrisome because employing dedicated snare removal teams is expensive, potentially limiting investment in other strategies to reduce snaring pressure.

Here, we used 11 years of patrol data from two contiguous protected areas in central Viet Nam to assess the impact of snare removal on snaring levels over time. Like many protected areas in Southeast Asia, the reserves are under high levels of snaring pressure, and snare removal has been the main strategy to counteract this threat. Within this context, we seek to understand how snare removal fits within a larger framework to counter snaring, including measures that prevent snares from being set; we refer to these approaches as “reactive” and “proactive”, respectively, even though we acknowledge that these two categories are not always clear cut, and reactive patrolling may also have proactive effects discouraging setting of snares (Moore et al, 2018; Dancer et al, 2022; results of this study). To understand the financial investment needed for snare removal, we also investigated the cost needed to implement these efforts, and possible sustainable opportunities to support this undertaking. We discuss our findings within the wider context of a holistic framework for the long-term objective to stop snaring in tropical protected areas.

## Methods

### Study site

Snare data were gathered by ranger patrols in the Hue Saola Nature Reserve (15,622 ha) and Quang Nam Saola Nature reserve (15,965 ha), located in central Viet Nam (Figure S1). The two reserves form one contiguous forest area across a provincial boundary. They are characterized by closed canopy broadleaf tropical rainforest, rugged terrain, and elevations ranging from 90 to 1450 m. A highway passes through the eastern section of the reserves.

Hunting is illegal in the protected areas, but nonetheless commonplace (Gray et al, 2014; Tilker et al, 2019). The main method of hunting is by setting wire snares, but other trap types – such as log-fall traps (MacMillan & Nguyen, 2014) – are also used found. In this study we analyzed all traps (wire snares and other traps) together, but because the majority of trapping is done by snaring (over 85% of traps were wire snares), we use the term “snares” throughout to align our phrasing to the commonly used and well referred term ‘snaring crisis’ for Southeast Asia (Belecky & Gray, 2020). Snaring in the two protected areas is commercially motivated, as it is across Viet Nam, (Belecky & Gray, 2020; Gray et al, 2021), with wildlife products supplying the high demand for bushmeat that exists throughout the country (Van Song, 2008). To counter this threat, ranger teams consisting of WWF Forest Guard members, often accompanied by a government Forest Protection Department representative, patrol the reserves with the primary mandate of finding and removing snares. The patrols started in 2011 and are ongoing. During the initial years of patrolling, costs were covered by WWF, with financing provided through international biodiversity conservation aid funds. In 2014, patrol costs began to be supplemented by a Payment for Forest Environmental Service (PFES) initiative by the Vietnamese government, and provided directly to the provincial management authorities. Some Forest Guard costs began to be covered by PFES funding starting in 2015, increasing to 15% in 2021 (WWF unpublished data).

### Collecting and processing patrol data

Ranger patrols were conducted in the contiguous Hue and Quang Nam Saola Nature Reserves by Forest Guard teams, coordinated by World Wide Fund for Nature (WWF), together with local Forest Protection Department staff. Patrols were conducted on foot, with temporary camps in the forest when needed, and lasted an average of 2.6 days, with 95% of patrols between 1 and 8 days in duration. Threat data was input into Management Information System (MIST) and Spatial Management and Reporting Tool (SMART) systems, and these data were curated and cleaned to create a database for analyses. We used patrol tracklog data to calculate monthly patrol effort in a grid of 200 x 200 meter cells as the percentage of each cell’s area surveyed. Tracklogs were buffered by 20 meters to calculate the total size of the buffered area within each cell. Data for snares were converted into detection/non detection matrices, with “0” representing no snares detected and “1” representing one or more snares detected in each 200 x 200 m cell. We then collapsed detection histories into monthly occasions. Snares that could not be assigned to a tracklog were removed from the analyses (Table S1). See Supporting Information for details.

### Description of the covariates

We considered the effect of five site covariates which we assumed could influence snare occupancy: village density, remoteness, elevation, terrain ruggedness (topographic ruggedness index), and topographic position index (TPI) (Figure S2). All covariates were calculated in QGIS 2.18.9 (QGIS Development Team, 2016) and R version v4.1.3 (R Core Team, 2022). Village density was calculated following Tilker et al. (2020), first creating a ground-truthed shapefile documenting villages around the study sites, then using a kernel density estimation in QGIS 2.18.9 (QGIS Development Team, 2016) to create a village density heatmap. Remoteness was calculated to provide the time required to reach every grid cell in the landscape from the nearest access points. It was assessed based on a 30-m resolution Shuttle Radar Topography Mission (SRTM) digital elevation model (Farr et al, 2007) by calculating time taken to traverse each pixel as a function of slope via Rees’ correction of Naismith’s rule (Rees, 2004), and then calculating cumulative cost (i.e., walking time) from major access points to each point in the landscape. Elevation was calculated from the 30-m SRTM digital elevation model (Farr et al, 2007). TPI was calculated from the 30-m SRTM using the t*errain* function in the R package *raster* v.3.3-7 (Hijmans et al, 2015). We did not find collinearity among covariates (|ρ| ≤ 0.45; see Table S2). See Supporting Information for more details.

### Model description

We used a multi-season occupancy model (Mackenzie et al, 2017) to assess the distribution of snares across both reserves. We implemented the model in a Bayesian framework in R using the *ubms* package v1.1.9005 (Kellner et al, 2022). We divided the survey period into semesters (the first and second half of each year) and used these as primary occasions. The second semester of 2012 and the first semester of 2013 were removed due to missing data.

The final model included the effect of survey effort as a covariate on detection probability and the five site covariates on occupancy probability; a random effect of year on the coefficients of all occupancy covariates and the occupancy intercept (allowing covariate coefficients and baseline occupancy to vary between years); and a random effect of year-semester on detection probability (allowing variation in detection probabilities, e.g. due to changes in staff). We found no evidence of spatial autocorrelation. All models converged, and showed adequate fit (see Supporting Information).

We used parameter estimates to predict occupancy probabilities for each year and raster cell. From these, we calculated the percentage of area occupied (PAO) as the percentage of cells predicted to be occupied by snares in each year for each posterior sample. The resulting distributions of PAO values were summarized to obtain mean and 95% Bayesian credible intervals (BCIs) of PAO.

We also used occupancy models to assess the influence of past patrolling effort on snare occupancy. Specifically, we fit separate single-season occupancy models to data from the second semester of 2016 and 2021, using the same covariate structure as described before, and adding as a covariate on occupancy the total snare patrolling effort summed over the preceding six semesters, calculated for each cell. We picked these two semesters as the approximate mid and endpoint of our time series, and the preceding time frame of six semesters because that was the longest preceding interval for which we had reliable patrol effort data for these two time points (see Supplementary Information).

### Financial investment

We estimated the costs to remove a single snare by the Forest Guards, with data provided by WWF (WWF internal and unpublished data), and the cost to set a snare by a local hunter, including opportunity cost based on the average daily salary for a worker, with cost estimates provided by consultation with experts familiar with the study area (see Supporting Information).

## Results

### Patrolling effort and snares removed

Between 2011 and 2021, the snare removal teams patrolled 3,040 days, covering 253,048 ha (Figure 1A). Patrol effort varied among and within years (Figure 1A; Figure S3). Overall, 99.2% of all 8,147 200 x 200 m cells received some level of effort within the study period (Figure S2). Patrol intensity was higher near major forest access points and in more accessible areas, and lower in more remote areas (Figure S3). Patrols removed 118,151 snares from the two reserves (Figure 1A), with more snares removed from the Hue Saola Nature Reserve compared to Quang Nam Saola Nature Reserve (Figure S4). The average patrol effort required to remove a snare across all years was 2.14 ha per trap collected. Average effort required to remove a snare increased across the study period from 1.3 ha per trap in 2011 to 2.62 ha per trap in 2021, with a peak of 4.14 ha per trap in 2019, indicating that over the course of the study period it took more effort to find a single snare (Figure S5).

**Figure 1:**
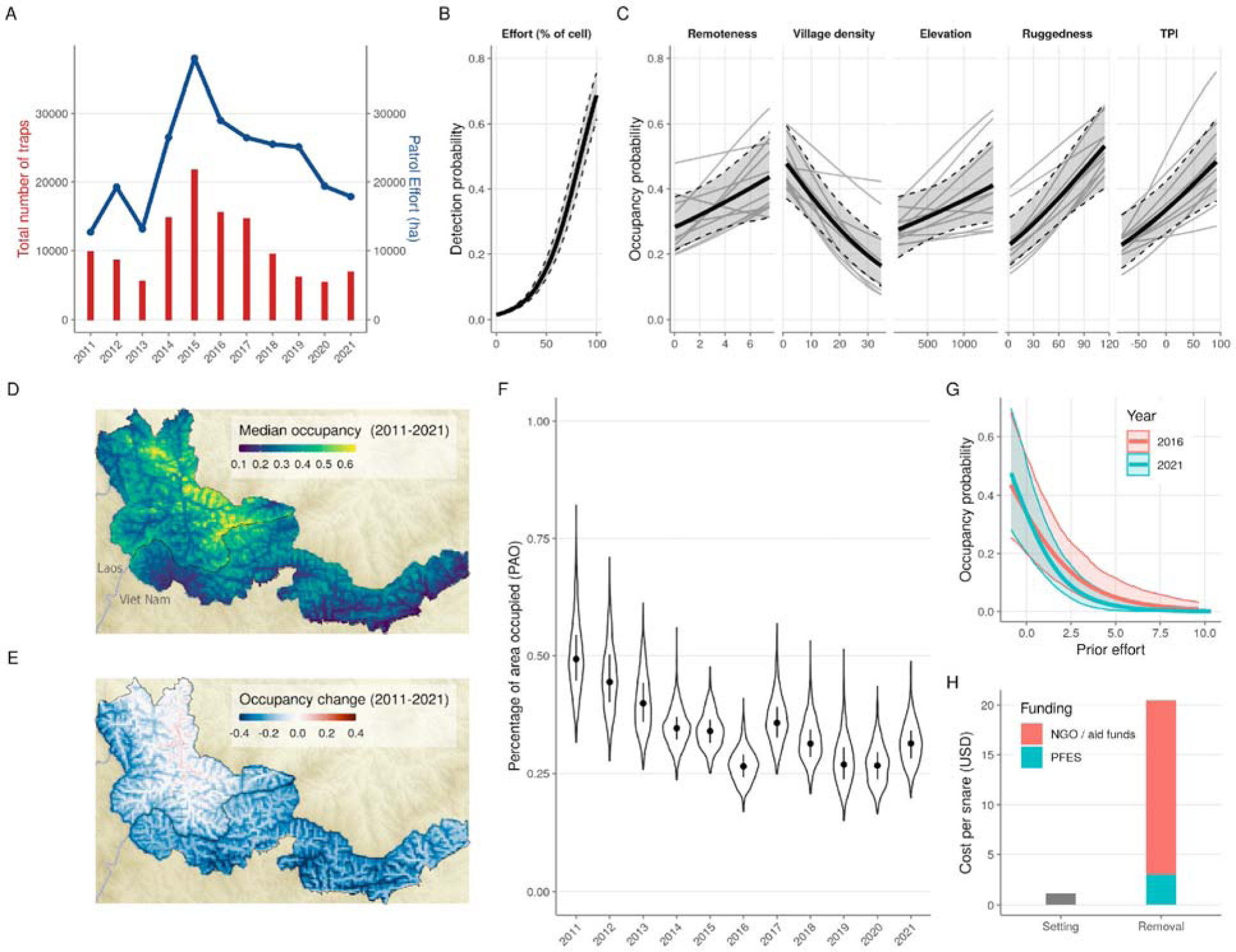
(A) Total number of snares removed by Forest Guards (blue line) and patrol effort in ha (red line) ; (B) response of snare detection probability to survey effort; (C) response of snare occupancy probability to covariates; thick black line shows mean effect for all years, gray shaded area show 95% confidence intervals (B and C), and light gray lines show mean effect for individual years (C only); (D) median snare occupancy probability for survey period (2011-2021); (E) median change in snare occupancy probability across the survey period; (F) Percentage of area occupied (PAO) values for individual years; black dot indicates median and line indicates 50% posterior mass; based on predictions from multi-season occupancy model (D - F); (G) response of snare occupancy to prior patrol effort for two time periods, 2016 and 2021; and (H) cost of setting and removing an individual snare in the study areas, with removal cost divided by Payment for Forest Ecosystem Services and NGO aid funds.

### Snare occupancy over time

The multi-season occupancy model showed that the mean snare percent area occupied (PAO) decreased from 0.500 (95% BCI [0.368, 0.666]) in 2011 to 0.314 (95% BCI [0.236, 0.408]) in 2021, resulting in a 36.9% (95% BCI [4.6, 59.0]) reduction from 2011 to 2021 (Figure 1C, E & 1F; Figure S6). The largest decline was 46.1% (95% BCI [21.8, 64.0]) between 2011 and 2016, after which PAO remained approximately stable (Figure 1F). The decline in snare PAO was slightly stronger in the Quang Nam Saola Nature Reserve (Figure 1D). Snare detection probability increased with the amount of effort, as measured by percent area covered per cell. (Figure 1B). Predicted snare occupancy increased with topographic position index (TPI), elevation, remoteness, and ruggedness, and decreased with village density (Figure 1C; Table S3). There was no strong correlation between initial snare occupancy and percent change in occupancy over time (Spearman rank correlation = - 0.35), indicating weak evidence that cells with initial higher snare occupancy showed larger declines across the study period (Figure S7).

Finally, snare occupancy probability decreased with increasing previous patrol effort (Figure 1G); the effect was significant (95% BCIs did not overlap 0) in 2016 (-0.48, 95% BCI [-0.77, -0.24]) and 2021 (-0.67, 95% BCI [-0.99, -0.40]), but had a larger magnitude in 2021.

### Financial investment

To achieve this level of snare reduction, it was necessary to invest approximately 220,000 USD annually (WWF unpublished data), resulting in an average cost of 20.5 USD per snare removed; this is more than an order of magnitude more expensive than the cost of setting a single snare (1.13 USD; Figure 1H).

## Discussion

### Costs and benefits of snare removal

We show that, with intensive and sustained snare removal, snaring levels can be significantly reduced within tropical protected areas. Snare removal is therefore an important component of strategies to counter the widespread snaring in Southeast Asia, if employed at the appropriate spatial and temporal scales. Overall snare occupancy declined steadily for the first six years of patrolling, and then appeared to plateau. This may indicate that, after an initial decline in snaring, subsequent reductions are progressively more difficult to secure. Furthermore, declines in snaring were stronger near the edges of the reserves and along access points (Figure 1e), perhaps because these areas were closer to major access points and therefore patrolled more frequently. Such a finding highlights the importance of increasing patrol effort in remote areas, especially since these are more likely to harbor conservation-priority species (Tilker et al, 2020). In this way, robust monitoring of patrol data has the potential to offer insights into patrolling strategies that can help inform adaptive management.

Snaring appears to be driven by a complex set of landscape factors in our study sites (Figure 1c). We found that snare occupancy was higher in more remote areas. Other studies have found both inverse (Plumptre et al, 2014; Kimanzi et al, 2015) and positive (Sarkar et al, 2022) relationships between snaring and remoteness. In our landscape, we posit that hunters are spending more time in remote areas with higher wildlife densities. Tilker et al (2020) found that wildlife occurrence increased with remoteness in the study site, and it is therefore likely that hunters are targeting remote areas that will yield greater return on investment, and spending less time in more accessible but depleted areas. It is also possible that hunters are actively avoiding ranger patrols, which are more frequent around more accessible areas; studies have shown that poachers may adapt their behavior to evade ranger interdiction (Ibbett et al, 2021). We consider that the positive relationship between snares and village density, with more snares further away from village clusters, reflects a similar pattern. Our finding that snaring increases with elevation, terrain ruggedness, and along ridgelines (higher TPI) may be related to higher defaunation in areas characterized by lower elevation or easier terrain, and to the fact that hunters set snares on ridgelines because they may be used by animals as trails (Ancrenaz et al, 2012), though further information would be needed to fully understand these drivers.

Our finding that prior patrolling decreased the probability of snare occupancy suggests that patrolling acts as a deterrent to future snaring. Other studies have also indicated the deterrence value of patrolling (Moore et al, 2028; Dancer et al, 2022). One possible explanation is that, with continued snare removal, hunters experience increasingly diminishing rates of return, and thus the cost-benefit ratio shifted so that it was less profitable to set snares in the study areas. In other parts of the world, the threat of arrest or fines might also serve as a deterrent; in our sites in central Viet Nam, however, arrests for snaring are rare, due in part to legal loopholes that do not penalize people for having wire materials inside protected areas (Belecky & Gray 2020; Gray et al, 2021), and in part because snaring is seldom treated as a serious forest crime. We note that it is possible that local reductions in snaring in our sites caused poaching to spill over to other protected areas which did not receive intensive patrol effort; similar patterns of displacement have been documented for illegal logging in tropical protected areas (Ford et al, 2020). Additional studies are needed to assess potential leakage and, if present, ways to counteract it.

Despite the success of demonstrably reducing snaring levels in these protected areas, the snaring levels that persist suggests that wildlife populations are still threatened by unsustainable offtake. Snaring has already contributed to the loss or functional extinction of many larger vertebrates from these areas (Tilker et al, 2019) – including the saola (*Pseudoryx nghetinhensis*), a Critically Endangered bovid that these reserves were established to protect – and it seems likely, based on other studies that have indicated that high levels of snaring can have severe negative impacts wildlife (Noss, 1998; Gray et al, 2018; Belecky & Gray, 2020), that remaining conservation-priority and snaring-sensitive species will continue to decline unless snaring pressure is reduced further. However, we note that because species show different abilities to recover from unsustainable harvest, largely due to life history characteristics (Cardillo et al, 2008), it is possible that populations of some mammals and ground-dwelling birds will stabilize or increase with this level of snare reduction. Long-term wildlife monitoring is needed to assess population dynamics and potential recovery for individual species.

### Perspectives on snaring in Southeast Asia

Reducing and maintaining decreased snaring levels in our study sites required long-term investment with considerable financial cost. The constant effort needed to reduce snaring levels may require conservation stakeholders to re-think how they approach site-based conservation, especially since snare-removal efforts are often funded by short-term conservation or development aid projects that last at best a few years. The financial investment necessary to attain such a level of snare reduction was substantial (>200,000 USD annually). With an estimated 13 million snares in the protected areas of Viet Nam, Laos, and Cambodia alone (Belecky & Gray, 2020), the cost to remove existing snares in Southeast Asian protected areas would likely sum to hundreds of millions of dollars. Moreover, the high cost of snare removal prompts questions about the long-term sustainability and scalability of this approach. It was only with a substantial and continual investment from an external NGO that such intensive levels of snare removal were possible in the two sites. Given the general lack of financial and personnel resources, capacity, and government support that characterizes most protected areas in the region (Graham et al, 2021), similar snare removal projects can likely only be implemented if NGO stakeholders work in collaboration with government protected area staff. One option to increase the financial sustainability of snare removal is to access sustainable financing schemes, such as Payment for Ecosystem Services (PES, PFES in Viet Nam), Reducing Emissions from Deforestation and forest Degradation (REDD+), or Carbon and Biodiversity offsets. The use of such incentive programs seems to be increasing: In 2020, more than 80% of protected areas in Viet Nam obtained PFES funds at an average of 323 USD per km^2^ (Emerton et al, 2021), which based on Decree No. 156/2018/ND-CP can be used for forest and biodiversity protection activities. However, these sustainable financing schemes are unlikely to be a shortcut to complete financing of snare removal operations; in the two reserves in our study, only 15% of the total costs could be covered by PFES in 2020 and 2021. Furthermore, funding alone will not guarantee effective replication of intensive snare removal in Southeast Asian protected areas. To guarantee transparency and accountability, protected area managers should set clear targets for threat reduction, and these metrics should be monitored. Our analytical workflow provides a streamlined approach to facilitate such robust analyses of patrol data in the future.

Our results suggest that snare removal alone may be insufficient to protect wildlife in Southeast Asian protected areas, especially for rare or snaring-sensitive species, since snaring levels remained high despite the demonstrated reductions. Although this is not unexpected, since snare removal does not address the fundamental drivers behind snaring, our results provide strong empirical evidence that a multifaceted approach is needed that combines short-term immediate responses within protected areas, such as snare removal, with long-term approaches in rural and urban areas (Figure 2). Such long-term responses could include demand reduction for illegal wildlife products (Shairp et al, 2016), strengthened legislation that penalizes the possession of wire material inside protected areas (Belecky & Gray, 2020), community engagement that promotes local guardianship (Viollaz et al, 2022), improved law enforcement coordination and capacity (Dudley et al, 2013), and engagement with local judicial authorities to ensure appropriate prosecution and sentencing (Nurse & Nurse, 2015). Simultaneously, actions which support inclusive governance of protected areas and nature-based and nature-positive development opportunities for buffer zone communities are required to offset potential losses of income, ensure equitable benefit sharing, and harmonize both economic advancement and biodiversity management and protection (Borrini et al, 2004; Andrade & Rhodes, 2012). Alternative livelihood opportunities may be needed in contexts where hunting is linked to subsistence or extreme poverty (van Vliet, 2011), though this is generally not the case in Viet Nam (Drury, 2011; Harrison et al, 2016), including in our study sites. Together, the activities needed for such a holistic approach will involve coordination among multiple government and non-governmental stakeholders and will require considerable financial investment.

**Figure 2:**
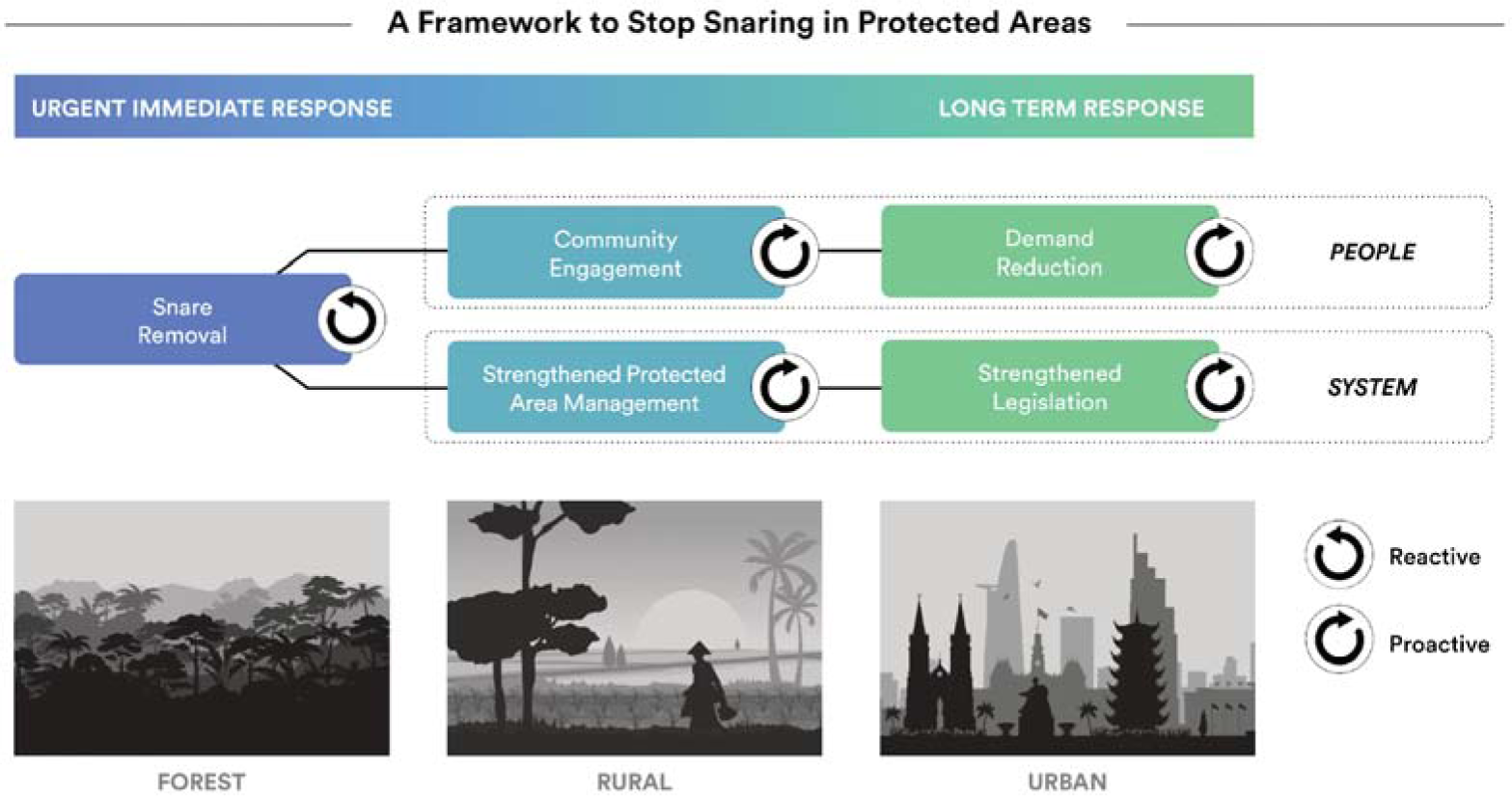
Conceptual framework to stop snaring in protected areas using a combination of reactive and proactive approaches across different environments and time scales.

Overall, our results are encouraging in that they show that sustained snare removal can be an effective method for reducing snaring. Therefore, intensive snare removal schemes should be expanded to other protected areas as a first and immediate response to safeguard highly threatened wildlife populations. For these populations, snare removal could buy crucial time needed to implement more proactive and long-term approaches, many of which could take time to become effective. Ideally, these efforts should be funded through sustainable financing mechanisms to allow for the long-term continuity necessary to suppress snaring. We acknowledge that, because the drivers behind snaring are so complex, the impact of sustained and intensive snare removal in other areas may differ from the results that we present here. Nonetheless, the high demand for wild meat in urban centers that is fueling snaring in central Viet Nam is present across Southeast Asia (Lee et al, 2014; Belecky & Gray, 2020), and we therefore consider the situation in our sites to be representative of the larger snaring problem in the region. Perhaps most importantly, our findings illustrate the importance of viewing snare removal as one component of a wider and multi-faceted conservation response that addresses these underlying drivers; to rely solely on snare removal to safeguard protected areas is unlikely to sufficiently address the threat. If we want to prevent the extinction of many of the region’s iconic megafaunal species, as well as to preserve the health of the tropical ecosystems that Southeast Asian societies depend upon, snare removal operations will need to be embedded within a more holistic and proactive framework that prevents snares from being set in the first place.

## Acknowledgements

We thank the staff of the WWF-CarBi project (phase I and phase II), the Thua Thien Hue Saola Nature Reserve, and the Quang Nam Saola Nature Reserve for supporting data collection, including through providing personnel for snare removal. The CarBi projects were and are still funded by the International Climate Initiative of the Federal Ministry for the Environment, Nature Conservation, Nuclear Safety and Consumer Protection (BMUV) through the German Development Bank (KfW). A.T. would like to thank Barney Long for productive discussions over the years on snare removal within the context of protected area management, Julie Louvrier for advice on study design and analysis, and Lucy Emerton for input on protected area financing in Viet Nam .

## References

Ancrenaz, M., Hearn, A. J., Ross, J., Sollmann, R., & Wilting, A. (2012). *Handbook for wildlife monitoring using camera-traps* . BBEC II Secretariat c/o Natural Resources Office, Kota Kinabalu, Sabah, Malaysia. https://www.hutan.org.my/wp-content/uploads/Reports/Other#20reports/Camera_trap_manual.pdf

Andrade, G. S., & Rhodes, J. R. (2012). Protected areas and local communities: an inevitable partnership toward successful conservation strategies? Ecology and Society, 17(4).

Barlow, J., França, F., Gardner, T.A., Hicks, C.C., Lennox, G.D., Berenguer, E., Castello, L., Economo, E.P., Ferreira, J., Guénard, B. & Graham, N. A. (2018). The future of hyperdiverse tropical ecosystems. Nature, 559(7715), 517–526.

Becker, M., McRobb, R., Watson, F., Droge, E., Kanyembo, B., Murdoch, J., & Kakumbi, C. (2013). Evaluating wire-snare poaching trends and the impacts of by-catch on elephants and large carnivores. Biological Conservation, 158, 26–36.

Belecky, M. & Gray, T.N.E, (2020). Silence of the Snares: Southeast Asia’s Snaring Crisis, WWF International. https://wwfeu.awsassets.panda.org/downloads/wwf_snaring_report_2020.pdf?364669/Silence-of-the-Snares

Bello, C., Galetti, M., Pizo, M.A., Magnago, L.F.S., Rocha, M.F., Lima, R.A., Peres, C.A., Ovaskainen, O. & Jordano (2015). Defaunation affects carbon storage in tropical forests. Science Advances, 1(11), e1501105.

Benítez-López, A., Alkemade, R., Schipper, A. M., Ingram, D. J., Verweij, P. A., Eikelboom, J. A. J., & Huijbregts, M. A. J. (2017). The impact of hunting on tropical mammal and bird populations. Science, 356(6334), 180–183.

Benítez-López, A., Santini, L., Schipper, A. M., Busana, M., & Huijbregts, M. A. (2019). Intact but empty forests? Patterns of hunting-induced mammal defaunation in the tropics. PLoS Biology, 17(5), e3000247.

Bi, S. G., Koné, I., Béné, J. C. K., Bitty, E. A., Yao, K. A., Kouassi, B. A., & Gaubert, P. (2017). Bushmeat hunting around a remnant coastal rainforest in Côte d’Ivoire. Oryx, 51(3), 418–427.

Bogoni, J. A., Peres, C. A., & Ferraz, K. M. (2020). Effects of mammal defaunation on natural ecosystem services and human well being throughout the entire Neotropical realm. Ecosystem Services, 45, 101173.

Borrini, G., Kothari, A., & Oviedo, G. (2004). Indigenous and local communities and protected areas: Towards equity and enhanced conservation: Guidance on policy and practice for co-managed protected areas and community conserved areas (No. 11). IUCN, Gland, Switzerland.

Campbell, K., Martyr, D., Risdianto, D., & Clemente, C. J. (2019). Two species, one snare: Analysing snare usage and the impacts of tiger poaching on a non-target species, the Malayan tapir. Biological Conservation, 231, 161–166.

Cardillo, M., Mace, G. M., Gittleman, J. L., Jones, K. E., Bielby, J., & Purvis, A. (2008). The predictability of extinction: biological and external correlates of decline in mammals. Proceedings of the Royal Society B: Biological Sciences, 275(1641), 1441–1448.

Coad, L., Watson, J.E., Geldmann, J., Burgess, N.D., Leverington, F., Hockings, M., Knights, K. & Di Marco, M. (2019). Widespread shortfalls in protected area resourcing undermine efforts to conserve biodiversity. Frontiers in Ecology and the Environment, 17(5), 259–264.

Dancer, A., Keane, A., Beale, C.M., Dobson, A.D., Amin, R., Freeman, R., Imong, I., Jones, K., Linkie, M., Long, B. & Collen, B. (2022). Evidence of deterrence from patrol data: Trialling application of a differenced-CPUE metric. Conservation Science and Practice, 4(8), e12746.

Drury, R. (2011). Hungry for success: urban consumer demand for wild animal products in Vietnam. Conservation and Society, 9(3), 247–257.

Dudley, N., Stolton, S., & Elliott, W. (2013). Wildlife crime poses unique challenges to protected areas. Parks, 19(1), 7–12.

Emerton, L., NguylZn, V. D., Bui, T. M. N., & Roth, M. (2021). Review of PA financial status in Viet Nam: ‘self-financing’ needs, options & ways forward. Report to Conservation, Sustainable Use of Forest Biodiversity & Ecosystem Services in Viet Nam. GIZ-Bio Project Phase II.

Fa, J. E., & Brown, D. (2009). Impacts of hunting on mammals in African tropical moist forests: a review and synthesis. Mammal Review, 39(4), 231–264.

Ford, S. A., Jepsen, M. R., Kingston, N., Lewis, E., Brooks, T. M., MacSharry, B., & Mertz, O. (2020). Deforestation leakage undermines conservation value of tropical and subtropical forest protected areas. Global Ecology and Biogeography, 29(11), 2014–2024.

Figel, J. J., Safriansyah, R., Baabud, S. F., & Herman, Z. (2023). Snaring in a stronghold: Poaching and bycatch of critically endangered tigers in northern Sumatra, Indonesia. Biological Conservation, 286, 110274

Graham, V., Geldmann, J., Adams, V. M., Grech, A., Deinet, S., & Chang, H. C. (2021). Management resourcing and government transparency are key drivers of biodiversity outcomes in Southeast Asian protected areas. Biological Conservation, 253, 108875.

Gray, T. N., Nguyen Quang, H. A., & Nguyen Van, T. (2014). Bayesian occupancy monitoring for Annamite endemic biodiversity in central Vietnam. Biodiversity and Conservation, 23(6), 1541–1550.

Gray, T.N., Hughes, A.C., Laurance, W.F., Long, B., Lynam, A.J., O’Kelly, H., Ripple, W.J., Seng, T., Scotson, L. & Wilkinson, N. M. (2018). The wildlife snaring crisis: an insidious and pervasive threat to biodiversity in Southeast Asia. Biodiversity and Conservation, 27, 1031–1037.

Gray, T.N., Belecky, M., O’Kelly, H.J., Rao, M., Roberts, O., Tilker, A., Signs, M. & Yoganand, K. (2021). Understanding and solving the South-East Asian snaring crisis. The Ecological Citizen, 4, 129–141.

Groenenberg, M., Crouthers, R., Yoganand, K., Banet-Eugene, S., Bun, S., Muth, S., & Gray, T. N. E. (2023). Snaring devastates terrestrial ungulates whilst sparing arboreal primates in Cambodia’s Eastern Plains Landscape. Biological Conservation, 284, 110195.

Harrison, R.D., Sreekar, R., Brodie, J.F., Brook, S., Luskin, M., O’Kelly, H., Rao, M., Scheffers, B. & Velho, N. (2016). Impacts of hunting on tropical forests in Southeast Asia. Conservation Biology, 30(5), 972–981.

Krause, H., Keane, A., Dobson, A. D., Griffin, O., Travers, H., & Milner-Gulland, E. J. (2021). Estimating hunting prevalence and reliance on wild meat in Cambodia’s Eastern Plains. Oryx, 55(6), 878–888.

Jachmann, H. (2008). Illegal wildlife use and protected area management in Ghana. Biological Conservation, 141(7), 1906–1918.

Kellner, K. F., Fowler, N. L., Petroelje, T. R., Kautz, T. M., Beyer Jr, D. E., & Belant, J. L. (2022). ubms: An R package for fitting hierarchical occupancy and N-mixture abundance models in a Bayesian framework. Methods in Ecology and Evolution, 13(3), 577–584.

Krause, T., & Tilker, A. (2022). How the loss of forest fauna undermines the achievement of the SDGs. Ambio, 51(1), 103–113.

Lacher Jr, T.E., Davidson, A.D., Fleming, T.H., Gómez-Ruiz, E.P., McCracken, G.F., Owen-Smith, N., Peres, C.A. and Vander Wall, S. B. (2019). The functional roles of mammals in ecosystems. Journal of Mammalogy, 100(3), 942–964.

Laurance, W.F., Carolina Useche, D., Rendeiro, J., Kalka, M., Bradshaw, C.J., Sloan, S.P., Laurance, S.G., Campbell, M., Abernethy, K., Alvarez, P. & Arroyo-Rodriguez, V. (2012). Averting biodiversity collapse in tropical forest protected areas. Nature, 489(7415), 290–294.

Lee, T. M., Sigouin, A., Pinedo-Vasquez, M., & Nasi, R. (2014). The harvest of wildlife for bushmeat and traditional medicine in East, South and Southeast Asia: Current knowledge base, challenges, opportunities and areas for future research. Occasional Paper 115. Bogor, Indonesia: CIFOR.

Linkie, M., Martyr, D. J., Harihar, A., Risdianto, D., Nugraha, R. T., Leader-Williams, N., & Wong, W. M. (2015). Safeguarding Sumatran tigers: evaluating effectiveness of law enforcement patrols and local informant networks. Journal of Applied Ecology, 52(4), 851–860.

MacKenzie, D. I., Nichols, J. D., Royle, J. A., Pollock, K. H., Bailey, L. L., & Hines, J. E. (2017). Occupancy estimation and modeling: inferring patterns and dynamics of species occurrence. Elsevier.

MacMillan, D. C., & Nguyen, Q. A. (2014). Factors influencing the illegal harvest of wildlife by trapping and snaring among the Katu ethnic group in Vietnam. Oryx, 48(2), 304–312.

Moore, J. F., Mulindahabi, F., Masozera, M. K., Nichols, J. D., Hines, J. E., Turikunkiko, E., & Oli, M. K. (2018). Are ranger patrols effective in reducing poaching-related threats within protected areas? Journal of Applied Ecology, 55(1), 99–107.

Noss, A. J. (1998). The impacts of cable snare hunting on wildlife populations in the forests of the Central African Republic. Conservation Biology, 12(2), 390–398.

Nurse, A., & Nurse, A. (2015). “National wildlife legislation and law enforcement policies.” In Policing wildlife: Perspectives on the enforcement of wildlife legislation, *pp.* 63-82.

Osuri, A.M., Ratnam, J., Varma, V., Alvarez-Loayza, P., Hurtado Astaiza, J., Bradford, M., Fletcher, C., Ndoundou-Hockemba, M., Jansen, P.A., Kenfack, D., Sankaran, M. (2016). Contrasting effects of defaunation on aboveground carbon storage across the global tropics. Nature Communications, 7(1), 1–7.

Plumptre, A.J., Fuller, R.A., Rwetsiba, A., Wanyama, F., Kujirakwinja, D., Driciru, M., Nangendo, G., Watson, J.E. and Possingham, H.P. (2014). Efficiently targeting resources to deter illegal activities in protected areas. Journal of Applied Ecology, 51(3), 714–725

QGIS Development Team. (2016). QGIS Geographic Information System. Open Source Geospatial Foundation. URL http://qgis.org

R Core Team (2022). R: A language and environment for statistical computing. R Foundation for Statistical Computing, Vienna, Austria. URL https://www.R-project.org/.

Sarkar, D, Bortolamiol, S, Jan F. Gogarten, Joel Hartter, Rong Hou, Wilson Kagoro, Patrick Omeja, Charles Tumwesigye, & Chapman, C. A. (2022). Exploring multiple dimensions of conservation success: Long-term wildlife trends, anti-poaching efforts and revenue sharing in Kibale National Park, Uganda. Animal Conservation, 25(4), 532–549.

Shairp, R., Veríssimo, D., Fraser, I., Challender, D., & MacMillan, D. (2016). Understanding urban demand for wild meat in Vietnam: implications for conservation actions. PloS One, 11(1), e0134787.

Tilker, A., Abrams, J.F., Mohamed, A., Nguyen, A., Wong, S.T., Sollmann, R., Niedballa, J., Bhagwat, T., Gray, T.N., Rawson, B.M., Guegan, F., Kissing, J., Wegmann, M., & Wilting, A. (2019). Habitat degradation and indiscriminate hunting differentially impact faunal communities in the Southeast Asian tropical biodiversity hotspot. Communications Biology, 2(1), 1–11.

Tilker, A., Abrams, J.F., Nguyen, A., Hörig, L., Axtner, J., Louvrier, J., Rawson, B.M., Nguyen, H.A.Q., Guegan, F., Nguyen, T.V., Le, M., Sollmann, R., & Wilting, A. (2020). Identifying conservation priorities in a defaunated tropical biodiversity hotspot. Diversity and Distributions, 26(4), 426–440.

van Vliet, N. (2011). Livelihood alternatives for the unsustainable use of bushmeat. Report prepared for the CBD Bushmeat Liaison Group. Technical Series No. 60, Montreal, Secretariat of the Convention on Biological Diversity.

Viollaz, J., Rizzolo, J. B., Long, B., Trung, C. T., Kempinski, J., Rawson, B. J., Reynald, D., Quang,. X. Q., Hien, N. N., Dung, C. T., Huyen, H. T., Dung, N. T. T., & Gore, M. L. (2022). Potential for informal guardianship in community-based wildlife crime prevention: Insights from Vietnam. Nature Conservation, 48.

Van Song, N. (2008). Wildlife trading in Vietnam: situation, causes, and solutions. The Journal of Environment & Development, 17(2), 145–165.

Watson, F., Becker, M. S., McRobb, R., & Kanyembo, B. (2013). Spatial patterns of wire-snare poaching: Implications for community conservation in buffer zones around National Parks. Biological Conservation, 168, 1–9.

Watson, J. E., Dudley, N., Segan, D. B., & Hockings, M. (2014). The performance and potential of protected areas. Nature, 515(7525), 67–73.

Wilkie, D. S., & Carpenter, J. F. (1999). Bushmeat hunting in the Congo Basin: an assessment of impacts and options for mitigation. Biodiversity & Conservation, 8, 927–955.

Young, H.S., Dirzo, R., Helgen, K.M., McCauley, D.J., Billeter, S.A., Kosoy, M.Y., Osikowicz, L.M., Salkeld, D.J., Young, T.P, & Dittmar, K. (2014). Declines in large wildlife increase landscape-level prevalence of rodent-borne disease in Africa. Proceedings of the National Academy of Sciences, 111(19), 7036–7041.

