## Supplementary information. for "Addressing the Southeast Asian snaring crisis: impact of 11 years of snare removal in a biodiversity hotspot"

### Supporting information

*Collection and processing of MIST and SMART data*

Ranger teams used two platforms to store threat data: Management Information System (MIST) and Spatial Management and Reporting Tool (SMART). MIST data started in February 2011 for the Hue Saola Nature Reserve (available until September 2012) and in February 2012 (available until November 2012) for the Quang Nam Saola Nature Reserve. No data was available from December 2012 until July 2013, as the data was lost during the transition from MIST to SMART. SMART data were available from May 2013. The last data included in the analysis were recorded in December 2021.

To determine the patrolling effort and create a uniform database for subsequent analyses we first needed to curate, clean, and organize MIST/SMART patrol data. We first exported the data from MIST/SMART into .csv and .gpx files. We then imported the MIST/SMART combined data for the two protected areas into R version v4.1.3 (R Core Team, 2022) and cleaned the patrol tracklog data by establishing a consistent coordinate system (WGS84/UTM zone 48N) and time zone and removing duplicate tracks. Data outside the study area were removed. Tracklogs were split into independent segments if consecutive track points were located more than 1000 meters from previous points to avoid connecting disjunct sections of GPS tracks.

Tracklogs were buffered by 20 meters to calculate the total size of the buffered area within each cell per month. A 20 meter buffer was chosen as the likely area that can be effectively searched by a team in the tropical rainforest terrain of the two reserves, based on conversations with Forest Guards and other people familiar with the area.

We then calculated the dates each cell was visited and percent area covered. We buffered roads by 30m and removed this area from the total area covered. Finally, we calculated a “fraction monthly coverage" value by dividing the patrolled area per cell and month by the size of the grid cell. Detections that could not be matched with effort were removed (5,168 total records; Table S1).

*Model structure*

The final model structure for occupancy probability ($\psi$) and detection probability ($p$) at cell *i* in year-semester *j* was:

$$logit(\psi_{ij}) = \alpha_{year[j]} + \beta_{year[j]}X_i$$

$$logit(p_{ij}) = \gamma_j + \delta \text{effort}_{ij}$$

where $\alpha$ and $\gamma$ are random (by year, and year-semester, respectively) intercepts, β are random year-specific coefficients associated with the five above described covariates X, and $\delta$ is the fixed effect of effort.

*Covariates*

Village density serves as a potential predictor for hunting pressure at the local scale (Koerner et al, 2017). To calculate village density, we followed the process outlined in Tilker et al. (2020). First, we created a ground-truthed point shapefile layer documenting local villages around the study sites, then we created a kernel density estimation in QGIS 2.18.9 (QGIS Development Team, 2016) using the village shapefile as the input point layer with the default quartic kernel decay function and a radius of 15 km. The radius was chosen so that the entire study landscape was covered in the final covariate layer. Observations in the field indicate that all parts of the study landscape, even the remote areas, were subject to some level of hunting pressure. The village density covariate is unitless, with larger values indicating areas with a higher density of villages in their surroundings.

The remoteness covariate measures walking time from access points to each point in the landscape and thus also serves as a potential determinant for hunting pressure. Starting points were created by digitizing major roads in and around our study site for 2014 (the approximate midpoint of the overall study period), using the *timelapse* function in Google Earth and converting these into points with a 100-meter spacing. Walking time from the starting points to each point in the landscape was calculated using the "cumulativeCost" function in Google Earth Engine (Gorelick et al, 2017). Cost to traverse a pixel was calculated from slope based on a 30-m resolution Shuttle Radar Topography Mission (SRTM) digital elevation model (Farr et al, 2007) via Rees' correction of Naismith's rule (Rees, 2004) and corrected for off-road travel with a factor of 1.67.

Elevation was calculated from the 30-m SRTM digital elevation model for each 200x200 m cell. Both ruggedness and TPI are measures of terrain complexity derived from elevation and related to accessibility. Previous work has shown that both terrain measures can influence hunter accessibility and animal occurrence (Estes et al, 2011; de la Torre et al, 2018). We used the t*errain* function in the R package *raster* v.3.3-7 (Hijmans et al, 2015) to calculate ruggedness and TPI from the 30-m SRTM digital elevation model.

*Testing for spatial autocorrelation, chain convergence, and model fit*

To assess potential spatial autocorrelation, we also calculated Moran's I of the model residuals following Moore & Swihart (2005) for distance categories 0-600m to 0-5000m. These trials showed that spatial autocorrelation in model residuals was generally low (0-0.06) and below the thresholds commonly applied for using autocovariates (0.2 or 0.4, see Crase et al. [2012] or Schuster et al. [2014]). We therefore decided that spatial autocorrelation in our models is not severe and did not warrant inclusion of autocovariates in the final models.

We ran three parallel Markov chains with 5,000 iterations, of which we discarded 2,500 as warm-up. We assessed chain convergence using the Gelman-Rubin statistic with values close to 1 indicating convergence. Model fit was assessed using the MacKenzie-Bailey chi-square goodness of fit test for occupancy models (MacKenzie & Bailey, 2004) with 500 posterior draws and found to be adequate ($p = 0.766$).

*Calculating financial investment for setting and removing snares*

WWF Viet Nam provided estimates of their total costs in one year to implement the full-time Forest Guard snare removal operations (WWF unpublished and internal data). These costs include all operating costs, salaries, allowances, food, and equipment costs for the patrols. For calculating the total costs to remove a snare we divided the total number of removed traps by the total costs (approximate annual costs * 11 years of patrolling). To calculate the total costs for setting a snare we used approximate costs for the wire used to make the snares (0.20 USD per snare), the food costs per day (1 USD per day), and the opportunity costs (average daily salary of a worker 13 USD) for the hunters and the estimated number of traps set per day (15).

Opportunity cost (USD per snare) was calculated using the following equation:

$$Cost\ per\ snare\ = \frac{Cost_{missed\ salary} + \ Cost_{food}\ +\ (Cost_{wire}\ \cdot snares\ per\ day)}{snares\ per\ day}$$

**Figures**

**Figure S1**

Covariates used in the final occupancy analysis.

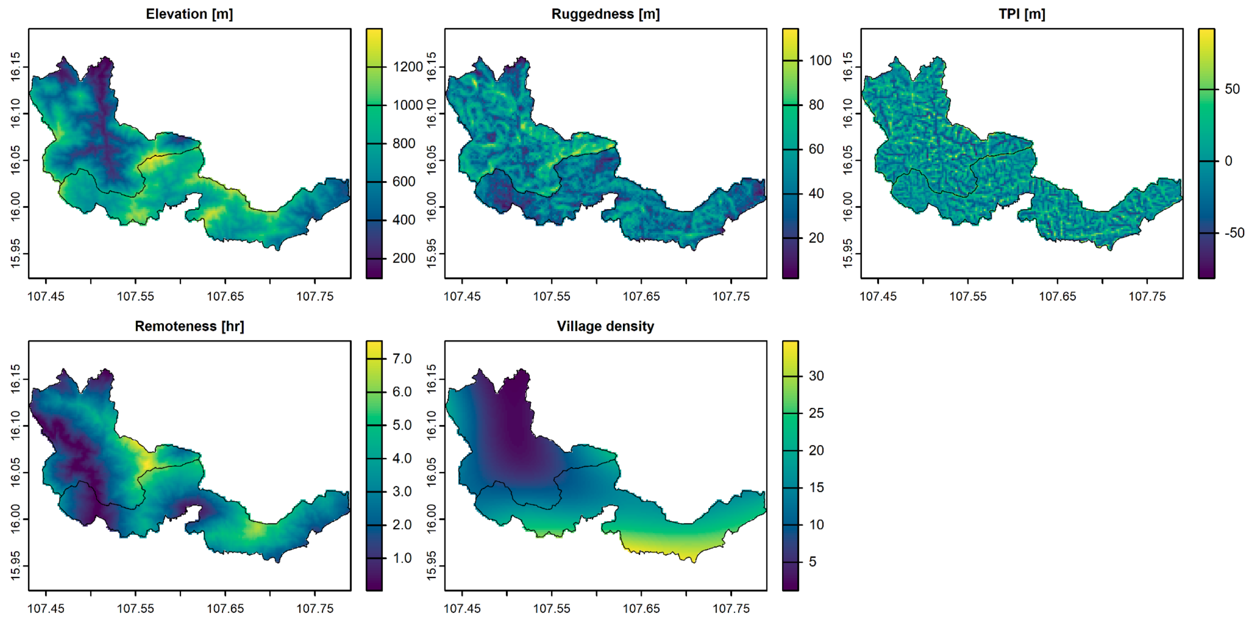


**Figure S2**

Figure showing patrol effort in the two reserves over the course of the study period.

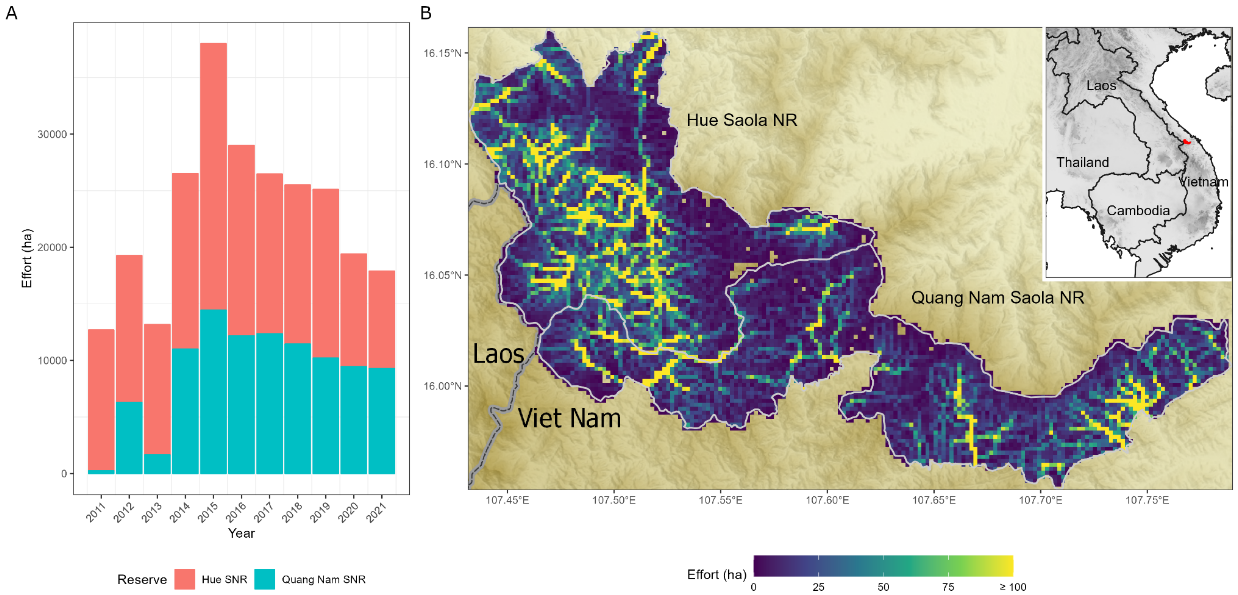


**Figure S3**

Figure showing cumulative snares recorded in the two reserves.

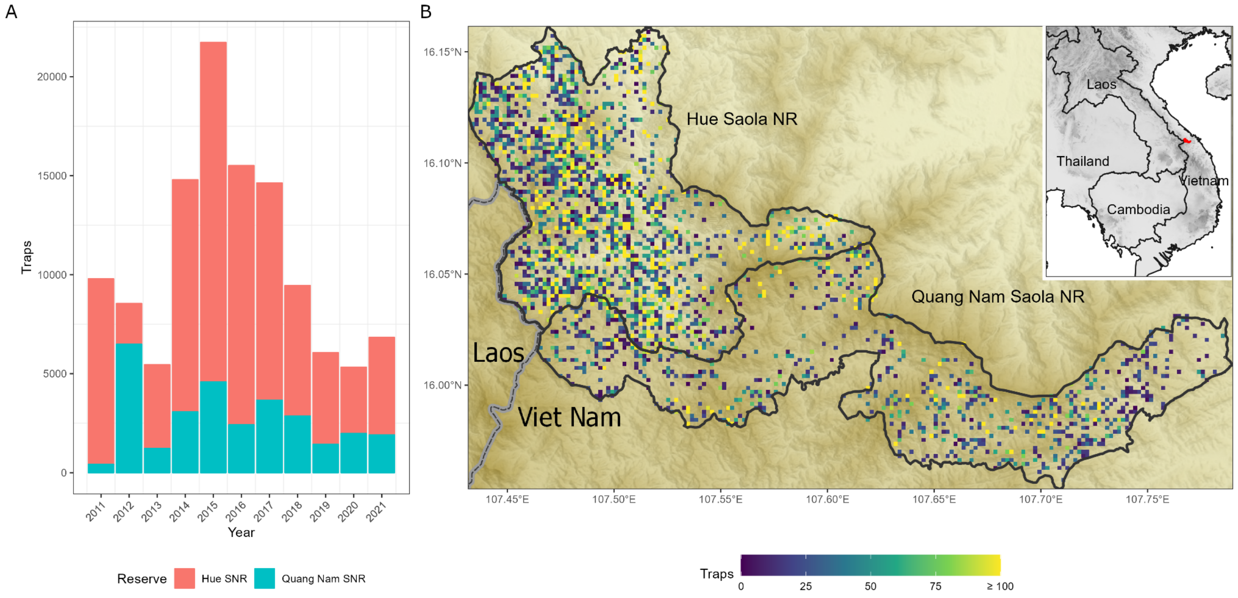


### Figure S4

Change in predicted snare PAO over time shown by year.

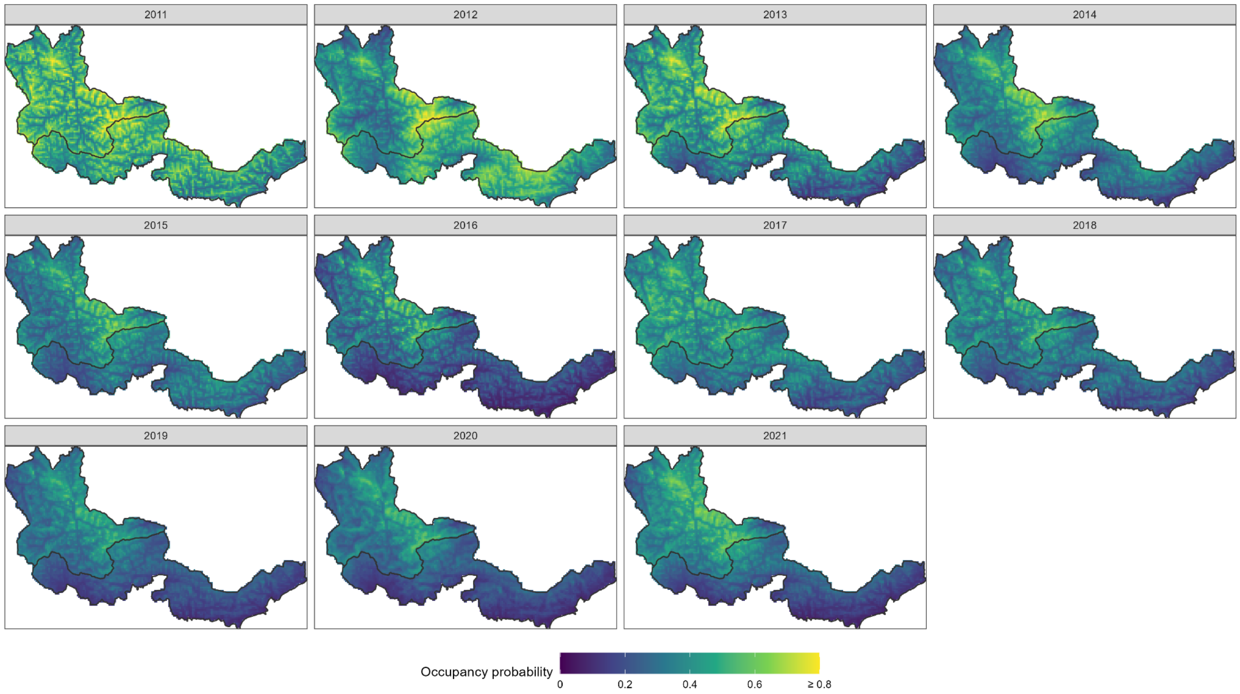

#### Tables

##### Table S1

Table showing number of records removed from analyses because they could not be matched to a given tracklog, divided by year and reserve.

| Year | Hue | QN |
|---|---|---|
| 2011 | 9375 | 420 |
| 2012 | 2060 | 6485 |
| 2013 | 4234 | 1220 |
| 2014 | 11725 | 3072 |
| 2015 | 17146 | 4585 |
| 2016 | 13095 | 2417 |
| 2017 | 10971 | 3665 |
| 2018 | 6590 | 2866 |

##### Table S2

Model results table (parameter estimates, SD, 95% BCI).

Occupancy

| | mean | sd | 2.50% | 97.50% |
|---|---|---|---|---|
| (Intercept) | -0.682 | 0.182 | -1.020 | -0.307 |
| elevation | 0.121 | 0.068 | -0.001 | 0.273 |
| village_density | -0.376 | 0.083 | -0.539 | -0.208 |
| remoteness | 0.143 | 0.065 | 0.014 | 0.271 |
| ruggedness | 0.200 | 0.046 | 0.110 | 0.292 |
| tpi | 0.167 | 0.050 | 0.073 | 0.269 |
| sigma [1\|year] | 0.460 | 0.168 | 0.203 | 0.845 |
| sigma [elevation\|year] | 0.146 | 0.077 | 0.025 | 0.325 |
| sigma [village_density\|year] | 0.226 | 0.082 | 0.091 | 0.417 |

| | | | | |
|---|---|---|---|---|
| sigma [remoteness\|year] | 0.166 | 0.068 | 0.056 | 0.324 |
| sigma [ruggedness\|year] | 0.089 | 0.059 | 0.008 | 0.225 |
| sigma [tpi\|year] | 0.117 | 0.063 | 0.016 | 0.262 |

Detection:

| | mean | sd | 2.50% | 97.50% |
|---|---|---|---|---|
| (Intercept) | -3.154 | 0.102 | -3.360 | -2.956 |
| effort | 0.826 | 0.023 | 0.780 | 0.872 |
| sigma [1\|year_semester ] | 0.327 | 0.081 | 0.197 | 0.507 |
